# Study of the variability of Pterosaur tracks from the Jurassic and Cretaceous of Spain using geometric morphometrics: a new insight on their ichnotaxonomy and anatomical implications

**DOI:** 10.1101/2025.08.05.668192

**Authors:** I. Jones, J. Marugán-Lobón, J.J. Moratalla

**Affiliations:** Unidad de Paleontología, Depto. Biología, Universidad Autónoma de Madrid. 28049 Cantoblanco (Madrid), Spain; Centro Para a Integración en Paleobiología, Universidad Autónoma de Madrid. 28049 Cantoblanco (Madrid), Spain; Instituto Geológico y Minero de España (IGME-CSIC), Museo Geominero. Ríos Rosas, 23. 28003 Madrid, Spain

**Keywords:** Paleoichnology, pterosaurs, *Pteraichnus*, Ichnotaxonomy, Geometric Morphometrics

## Abstract

Pterosaur tracks are relatively abundant in the Spanish fossil record and are especially localized in the Upper Jurassic of the Asturian coast and in the Lower Cretaceous of the Cameros Basin. All cases so far have been identified within the ichnogenus *Pteraichnus*. The Berriasian of the Cameros Basin is especially abundant in pterosaur ichnites, and, for this reason, the presence of seven ichnogenera had been proposed, about which, in general, there has not been a complete consensus among researchers. These ichnogenera were established according to normal procedure in paleoichnology, that is, with morphological characters, as well as considering several variables and meristic indexes. We provide a study on the variability of pterosaur handprints, focused on the Cameros Basin record, using shape analytical tools (geometric morphometrics). The results suggest that shape variability is continuous with respect to pterosaur handprint geometry. In consequence, there are no distinctive features that allow us to discriminate between them, even considering factors such as geological age or geographical origin. The shape of pterosaur manus prints remains very constant, suggesting a conservative evolution pattern of the module formed by the digits I, II and III, an important structure of the pterosaur arm for adequate quadrupedal locomotion on land.

## Introduction

Pterosaurs are a group of Mesozoic archosaurs, the first vertebrates that acquired the ability to fly (Unwin, 2005). However, they were also responsible for an important section of the impressive Mesozoic vertebrate fossil track record, dating from the Late Triassic and extending about 160My, until their extinction in the Late Cretaceous (Chatterjee & Templin, 2004). While the world fossil track record shows that dinosaurs have been the great producers of tracks in vertebrate history, giving rise to large areas sometimes with huge tracksites all over the world, pterosaurs shared many of such continental ecosystems with their dinosaur relatives, producing fossil tracks. However, the recognition of pterosaur ichnites as such was long ago shrouded in some confusion.

The starting of the consideration of pterosaur tracks probably began with Oppel (1862), who proposed that some tracks of the Late Jurassic Solhofen Limestone were attributable to the pterosaur *Ramphorhynchus*. This hypothesis was wrong as these tracks were after demonstrated to be of invertebrates (Malz, 1964; Wellnhofer, 1991), but it ignited interest in pterosaurs as potential trackmakers. Such debate was renewed by Stokes (1957) who reported some tracks from the Morrison Formation, and named them *Pteraichnus*, clearly assuming that they were in fact pterosaur tracks. However, other authors proposed the crocodile group as responsible for this ichnogenus (Padian & Olsen, 1984), an opinion that remained for years (Conrad et al., 1987; Unwin, 1989).

During these “intermediate” years, the peculiar tridactyl shape of the pterosaur manus prints led to consider them as possible aviform prints or even chelonian tracks (Moratalla, 1993). All these doubts were dispelled when *Pteraichnus* was finally demonstrated to be produced by a pterosaur and not by crocodiles or any other known group of reptiles (Lockley et al. (1995). Since then, a revolution in this field that allowed the recognition of many sites of pterosaur origin sharing many localities with dinosaur footprints and, on occasions, with crocodile and chelonian tracks, now much more clearly discriminated (Lockley et al., 2008). The relative abundance of pterosaur tracks has stressed the locomotor abilities of this group of archosaurs that, although dominating the Mesozoic skies, also wandered across the substrate.

The first pterosaur tracks in Spain were discovered at the site of Los Tormos (Santa Cruz de Yanguas, Soria) (Lower Cretaceous) and consisted basically of a trackway mainly formed by manus prints. These were of small size, with short digit marks, and sometimes scarcely marked, and arranged parallel to the axis of the trackway. In the mentioned trackway, some very shallow marks corresponding to the foot could be observed, poorly preserved and without a clear morphology. This trackway was at first potentially identified –as we have said before-with a chelonian (Moratalla, 1993) although it soon became evident that it was, in fact, a trackway produced by a pterosaur (Lockley et al., 1995). From this evidence, new fossil tracks of this type were discovered in the region, which highlighted the relative abundance of this type of ichnites in the Oncala Group (Berriasian) of the Cameros Basin.

From such report and other findings, it seems clear that the Spanish pterosaur track record is located mainly in two big areas: The Upper Jurassic of the Asturias coast (north of Spain) and the Lower Cretaceous of the Cameros Basin (Soria and La Rioja provinces). The Asturian coast presents a record mainly constituted by isolated foot and manus tracks, without the presence of trackways. Some specimens are very remarkable, are very well preserved and, sometimes, they show even very important details on skin structure and scale arrangement (García Ramos et al., 2000; Piñuela et al., 2007; Piñuela, 2015). In contrast, the Cameros Basin shows a remarkably different pterosaur record. Due to the different genetic, tectonic and sedimentary conditions of both areas, the pterosaur ichnites also present a very different preservation. Although, anatomically the details are not so precise, several Cameros pterosaur trackways have nevertheless been preserved in different sites that add data to better understand their mode of locomotion over the substrate (Moratalla et al., 1994; 2000; 2001; 2003; 2004; Moratalla, 2004; Moratalla & Hernán, 2009; Sánchez Hernández, 2009; Hernández-Medrano et al., 2014).

Interestingly, the ability to walk of pterosaurs today is unambiguously endorsed by a global record of tracks with more than 50 sites described, spanning four continents, and dating from the Middle Jurassic to the Early Cretaceous (Lockley et al., 2008). Thanks to the study of these trackways, it is known that pterosaurs had quadrupedal locomotion and were not bipedal and clumsy (Padian and Reyner, 1993) when it came to moving by land. This was probably caused by the location of the center of gravity near the shoulder girdle, which would make quadrupedal locomotion the usual way of moving over the substrate. This would imply the folding of the wings over the dorsum of the animal, giving rise to a digitigrade hand support, with the digits laterally directed while the feet made a plantigrade support with rather a forward orientation (Lockley et al., 1995; Chatterjee & Templin, 2004). Moreover, they probably could have had an upright gait and were quite agile, being able to run at relatively high speeds (Bennett, 1997) but, up to now, no pterosaur trackways showing a high-speed locomotion pattern have been reported.

Despite the relative abundance of pterosaur tracks, only three ichnogenera have been described to date: *Pteraichnus* (Stokes, 1957), *Purbeckopus* (Delair, 1963) and *Haenamichnus* (Hwang et al., 2002), suggesting a relatively low diversity, at least from a paleoichnological point of view. The purpose of this work is to analyze the shape variation of pterosaur tracks addressed as *Pteraichnus* using geometric morphometrics, as an attempt to clarify descriptive aspects of pterosaur ichnotaxonomy on which there is still no consensus among some authors (Moratalla et al., 2003).

## Material and methods

The sample encompasses N=180 pterosaur manus tracks from the Spanish Jurassic and Cretaceous fossil record scanned from the literature. Only manus tracks have been considered since their abundance is much higher than that of footprints (perhaps for functional aspects of locomotion and preservation bias). The sample comes from; 1) Cameros Basin; 2) Asturias coast and 3) Worldwide Fossil Track Record.

The Cameros Basin comprises more than 5,500 km^2^ of extension, forms the northernmost section of the Iberian rift complex that was active from the end of the Paleozoic to the Cretaceous, during the Aptian period (Mas et al., 2004). This Basin is located in the provinces of Soria, La Rioja and Burgos and has given rise to one of the regions with the largest number of fossil tracksites of the European continent (Moratalla, 1993; Moratalla & Hernán, 2009; Arribas & Pérez, 2000). The Cameros sediments are divided into five lithological groups (Tera, Oncala, Urbión, Enciso and Oliván) (Tischer, 1966 and Alonso et al., 1991). Of these five groups, three of them (Tera, Urbión and Oliván) have a mostly fluvial character while Oncala (Berriasiense) and Enciso (Upper Barremian-Lower Aptian) have a lacustrine character and, therefore, it is here where tracksites are much more abundant, with a vast majority of dinosaur tracks.

Pterosaur tracks are relatively abundant in the Oncala Group (Berriasian) with more than 50 sites having pterosaur traces, totaling up to about 2000 ichnites, most of them found in the Oncala group (Hernández-Medrano et al., 2017). On the contrary, from the Enciso Group (Upper Barremian-Lower Aptian), only pterosaur tracks have been found in Los Cayos locality (Cornago) (Moratalla & Hernán, 2009), and recently pterosaur traces have been found for the first time in the Tera group (Hernández-Medrano et al., 2017).

The ichnogenus *Pteraichnus* comprises ichnites of small to medium sizes, and it is defined by quadrupedal traces of hands and feet with the following characteristics according to Billon-Bruyat and Mazin (2003):

1. The manus prints are located on the same axis or laterally to the impressions of the feet.
2. Manus prints are imprinted equally or more deeply than footprints.
3. The footprints are elongated, slightly subtriangular, plantigrade, and tetradactyl, usually presenting claw impressions.
4. The manus tracks are tridactyl, digitigrade, and asymmetrical.
5. The digit I is printed in the anterior or anterolateral direction (it usually has a claw mark); the digit II shows an anterolateral to posterolateral direction (rarely presenting a claw mark), and the digit III shows a posterior orientation (exceptionally with a claw mark).

The manus digital marks increase respectively in length, with the longest being digit III. Up to 6 different ichnospecies have been described from the Cameros Basin (Table 1)

**Table 1.**
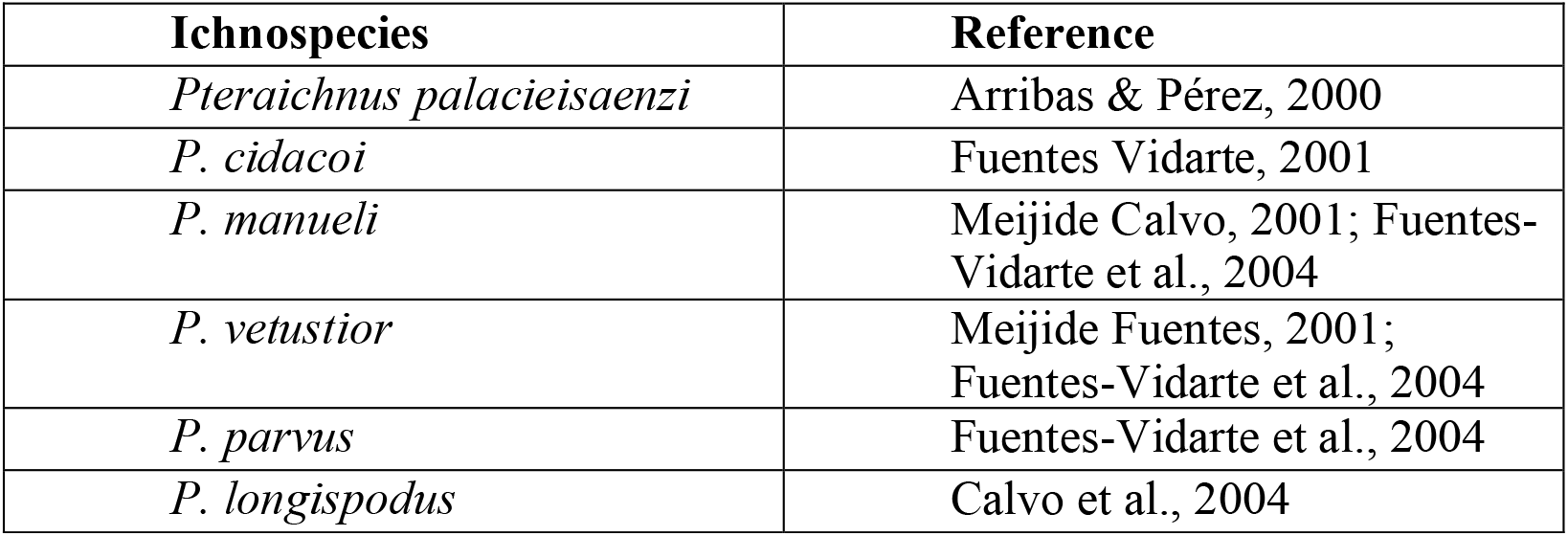
Ichnospecies described from the Cameros basin and their reference.

However, some ichnites have been taken as *numina dubia* due to the absence of a holotype exhibited in a museum or similar institution (Billon-Bruyat & Mazin, 2003), concluding that only *P. longipodus* and *P. parvus* are valid ichnospecies (Sánchez-Hernández et al., 2009). It is also relevant to mention the presence of another ichnospecies in the Spanish record, *P. cf. stokesi* (Arribas & Medrano, 2016), an ichnospecies described by Lockley et al. (1995) in the Sundance Formation of the Jurassic of Wyoming, USA.

The Asturias coast constitutes an important region with sediments of Late Jurassic (Kimmeridgian) in age (Table 2), specifically, from the Tazones Site in the Lastres Formation (Piñuela, 2015). The sedimentological context points to the existence of delta systems that allowed the formation of lagoons of variable sizes (Oloriz et al., 1988). The Tazones site is part of the Gijón-Villaviciosa Basin, specifically in the northern sector (Ramírez del Pozo, 1969), that is, all the materials of Jurassic origin that outcrop between Gijón and Ribadesella (Omeñaca et al., 2007). The tracks of this site have a fantastic conservation, even showing details such as interdigital hand membranes and even some tracks suggesting a process of take-off (Piñuela, 2015). Due to their hand morphology and the presence of tetradactyly footprints, they were classified within the ichnogenus *Pteraichnus* (Piñuela, 2015), coinciding with the pterosaur tracks from the Cameros Basin, as well as the rest of the ichnites included in this study.

**Table 2.** Ichnites used in the present study, according to the proposed ichnospecies, time, and site, each obtained from the corresponding bibliography. All come from the Early Cretaceous, except LR (1-8), considered to be the Upper Jurassic.

| Fingerprint ID | Ichnospecies | Deposit | Bibliography |
| --- | --- | --- | --- |
| HT-01 | *Pteraichnus cicadoi* | Oncala | (Arribas & Medrano, 2016) |
| HT-02 | *P. vetusitor* | Oncala | (Arribas & Medrano, 2016) |
| HT-03 | *P. longispodus* | Oncala | (Arribas & Medrano, 2016) |
| HT-04 | *P. manueli* | Oncala | (Arribas & Medrano, 2016) |
| HT-05 | *P. palaceisaenzi* | Oncala | (Arribas & Medrano, 2016) |
| HT-06 | *P. parvus* | Oncala | (Arribas & Medrano, 2016) |
| HT-07 | *P. cf stokesi* | Jurassic | (Arribas & Medrano, 2016) |
| BV-(1-72) | *P. sp* | Oncala | (Arribas & Pérez, 2000) |
| LC-(1-4) | *P. sp* | Enciso | (Moratalla & Hernán, 2009) |
| LM-(1-27) | *P. cf stokesi* | Oncala | (Arribas & Medrano, 2016) |
| LR-(1-8) | *P. sp* | Tera | (Hernández-Medrano et al., 2017) |
| EN-(1-14) | *P. palaceisanzei* | Oncala | (Arribas et al., 2014) |
| MJ-(1-42) | *P. sp* | Jurassic | (Piñuela, 2015) |
| V3-(1-2) | *P. parvus* | Oncala | (Fuentes et al, 2004) |
| BS-(1-4) | *P. longipodus* | Oncala | (Calvo et al., 2004) |

In addition to the ichnites from the Spanish record, 70 pterosaur tracks of the world fossil record have been added in a complementary way, which are referred to in Table 3. The aim was to expand the study sample to reinforce the consistency of the statistical results. These pterosaur tracks, attributed to the ichnogenus *Pteraichnus*, come from a great variety of worldwide sites (Table 3).

**Table 3.** Ichnites used in the comparative study, classified by their site, ichnogenera and age, each obtained from the corresponding bibliography.

| Fingerprint ID | Ichnospecies | Origin | Reference | Age |
| --- | --- | --- | --- | --- |
| NH-(1-11) | *Pteraichnus sp* | USA | (Lockley & Gillette, 1999) | Late Cretaceous (Maastrichtian) |
| WA-(1) | *P. sp* | China | (Xing et al, 2013) | Early Cretaceous (Aptian-Albian) |
| SF-(1-36) | *P. sp* | USA | (Mickelson et al, 2004) | Middle Jurassic (Callovian) |
| MF-(1-16) | *P. sp* | USA | (Lockley et al, 2001) | Late Jurassic |
| UF-(1-3) | *P. sp* | Korea | (Lockley et al, 1997) | Late Cretaceous |
| WQ-(1-3) | *P. sp* | Poland | (Pieńkowski & Niedźwiedzki, 2005) | Late Jurassic (Kimmeridgian) |

The track images were exported to the TPSdig2.64 computer software (Rohlf, 2015), through which the coordinates of 6 landmarks were digitized. The landmarks are corresponding Cartesian coordinates (in 2D in this case) of the tracks (i.e., between shapes), where the criterion is the hand biological homology in easily identifiable areas, such as the distal end of the digits, the hypexes between them, as well as the most posterior area of the track, considered as the projection of digit III (see Table 4 and Fig. 2).

**Table 4.** Landmarks used and their description.

| Landmark No. | Description |
| --- | --- |
| 1 | Distal End of Finger I |
| 2 | Hypex between fingers I and II |
| 3 | Distal End of Finger II |
| 4 | Hypex between fingers II and III |
| 5 | Distal End of Finger III |
| 6 | Projection of the finger II following its curvature |

**Figure 2.**
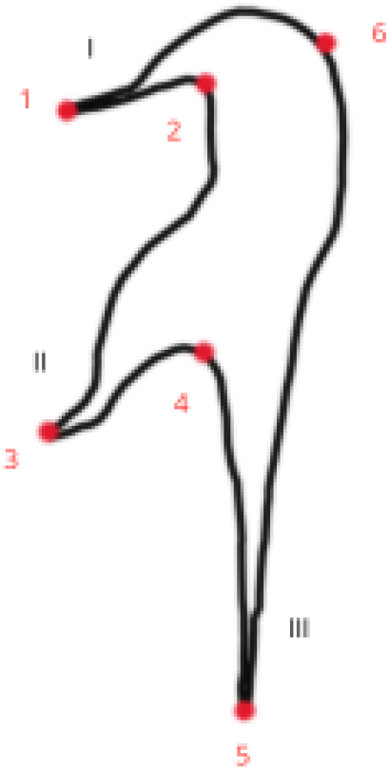
Diagram of the landmarks used in the study and the numbering of the fingers.

Once the data have been taken and correctly classified, the landmarks coordinates were exported to the statistical software for morphometric analysis, MorphoJ (Klingenberg, 2010) to obtain the Procrustes data (i.e., residuals; Rohlf & Slice, 1990, Fig. 3), after their superimposition under a Least Squares fit, namely, translating and rotating them to a common referent (the mean) following a least squares criterion and finally, rescaling them.

**Figure 3.**
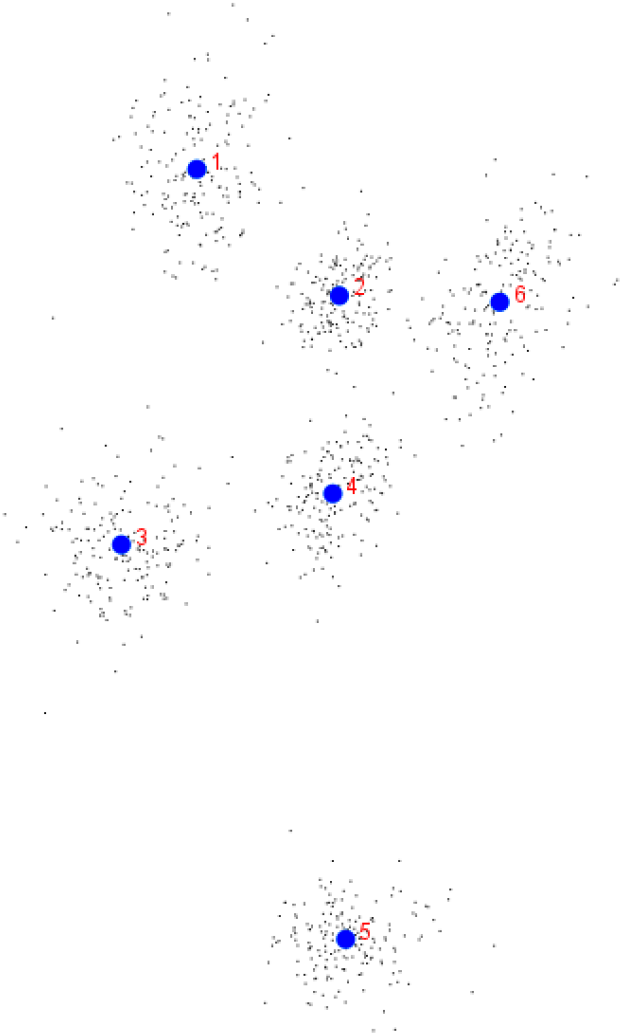
Residual values from the Procrustes Analysis (all overlapping configurations).

The analysis of the pterosaur tracks morphological (geometric) variation was summarized on a PCA Principal Component Analysis (PCA), a multivariate tool to reduce the dimensionality of the data, that is, it generates a few variables (linear combinations of the originals) that summarize the variance of the data (Jolicoeur, 1999). Discriminant function analyses (DFA) were performed on classifying pairings: 1) age (Jurassic-Cretaceous) and 2) registration (national-international) and analysis of canonical variables (CVA). The DFA analysis is based on equations that maximize the difference between groups, minimizing intragroup differences, and looking for the axes that best distinguish the groups defined in a priori (Aguirre & Prado, 2018). A cross-validation was carried out between the vectors (1000 permutations). The CVA is similar to the DFA, but it is used in variables with multiple groups (more than two) to assess if the groups present differences in their mean tendency, it must be noted that it can only be used as an approximation as the CVA will always separate groups when the number of variables is similar to the number of cases (Mitteroecker & Gunz, 2009).

Finally, track shape allometry was analyzed with a Multivariate Regression (Monteiro, 1999), taking as an independent variable the size of the centroid (the absolute size of the footprint that, from the landmarks, being computed as the square root of the sum of the squared differences (Zelditch, 2004). The significance is tested under a 10000 round permutation.

## Results

The 39% of the variance of the whole sample is explained by PC1, related to differences in the relative position of the fingers. On the other hand, PC2 explains 17% of the variance and stands for differences in the thickness of the digit marks (Fig. 4A). However, there is a high overlap (Fig. 4A) suggesting that there is no morphological difference depending on their age. Main shape differences affect the arrangement and the proportions of digits (PC1 and 2, respectively).

**Figure 4.**
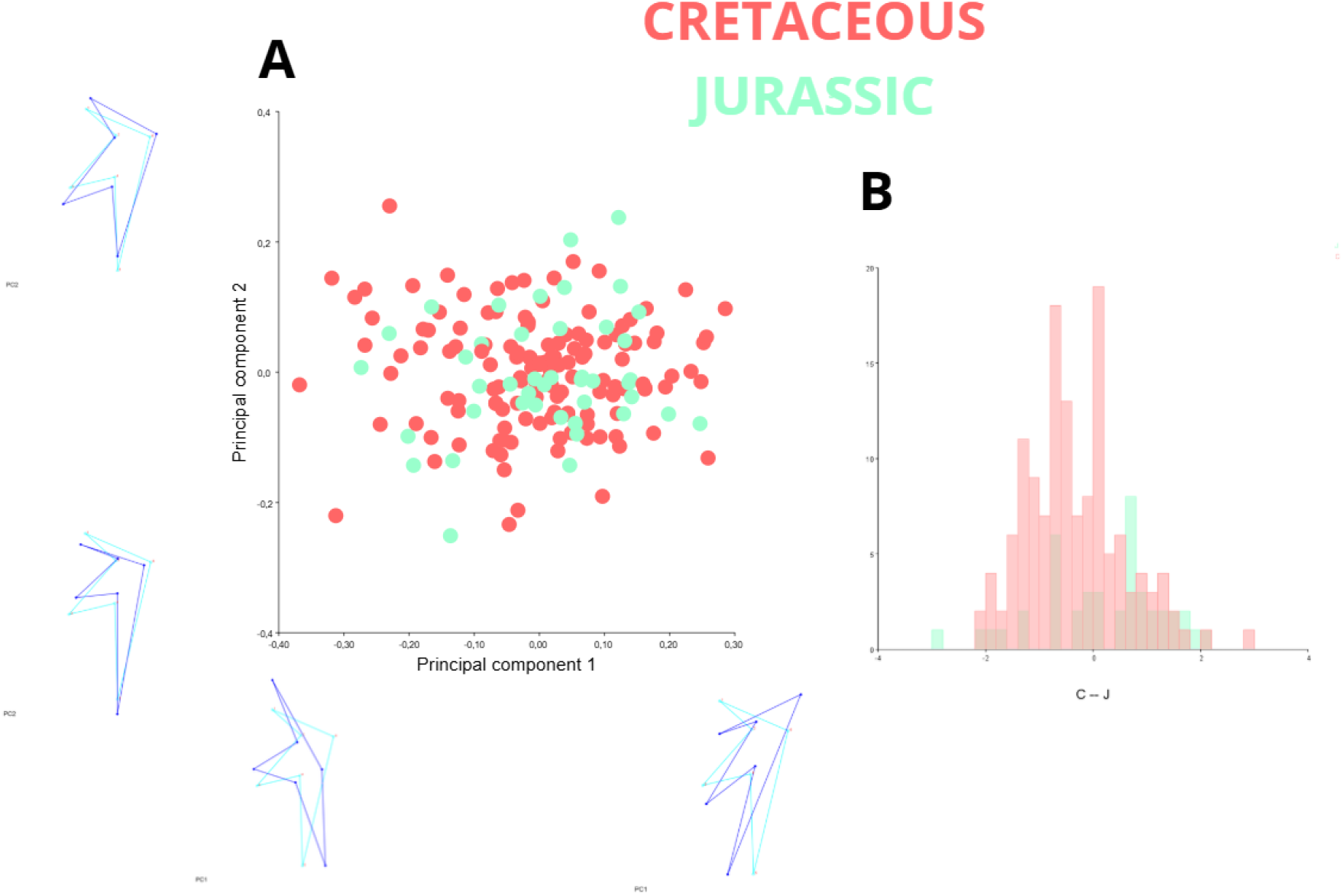
**A**, PCA results and variations in shape according to PC1 and PC2 compared to the mean. **B**, cross-validation of discriminant function analysis to find differences between the ichnites depending on their age (histogram).

After cross validating the discriminant function analysis (Fig. 4B), we found that separation between groups of observations is not statistically significant. Such a negative result is highly relevant since it clearly indicates that grouping shape data according to this criterion (qualitative identity) is impossible.

Comparing the data according to their origin (Spanish record-International record) the PCA (Fig. 5A) also shows that there is an overlapping of the data, implying that we cannot group the data according to their origin either. Again, cross-validation of the discriminant function (Fig. 5B) corroborates this.

**Figure 5.**
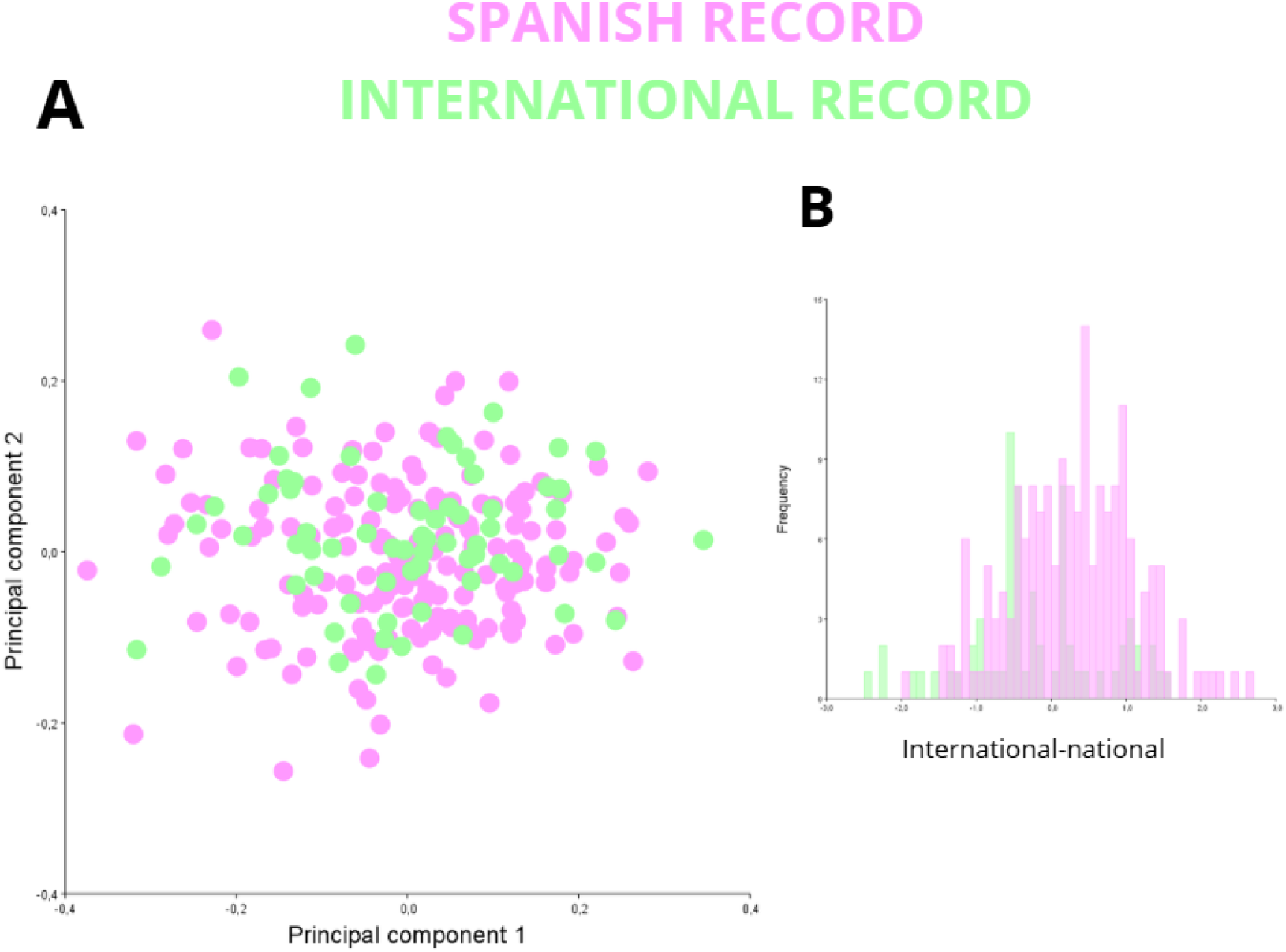
**A**, PCA according to the origin of the fingerprints. **B**, cross-validation of the discriminant function (histogram).

Regarding the pterosaur tracks from the Spanish record, the results of the PCA (Fig. 6A) show a high overlap between the ichnites of the different ichnospecies proposed and no clear groupings appear according to this criterion. The results of the Canonical Variable Analysis (CVA) (Fig. 6B) are significant since no type of ordering of these is observed, and they appear with a remarkably high degree of overlapping.

**Figure 6.**
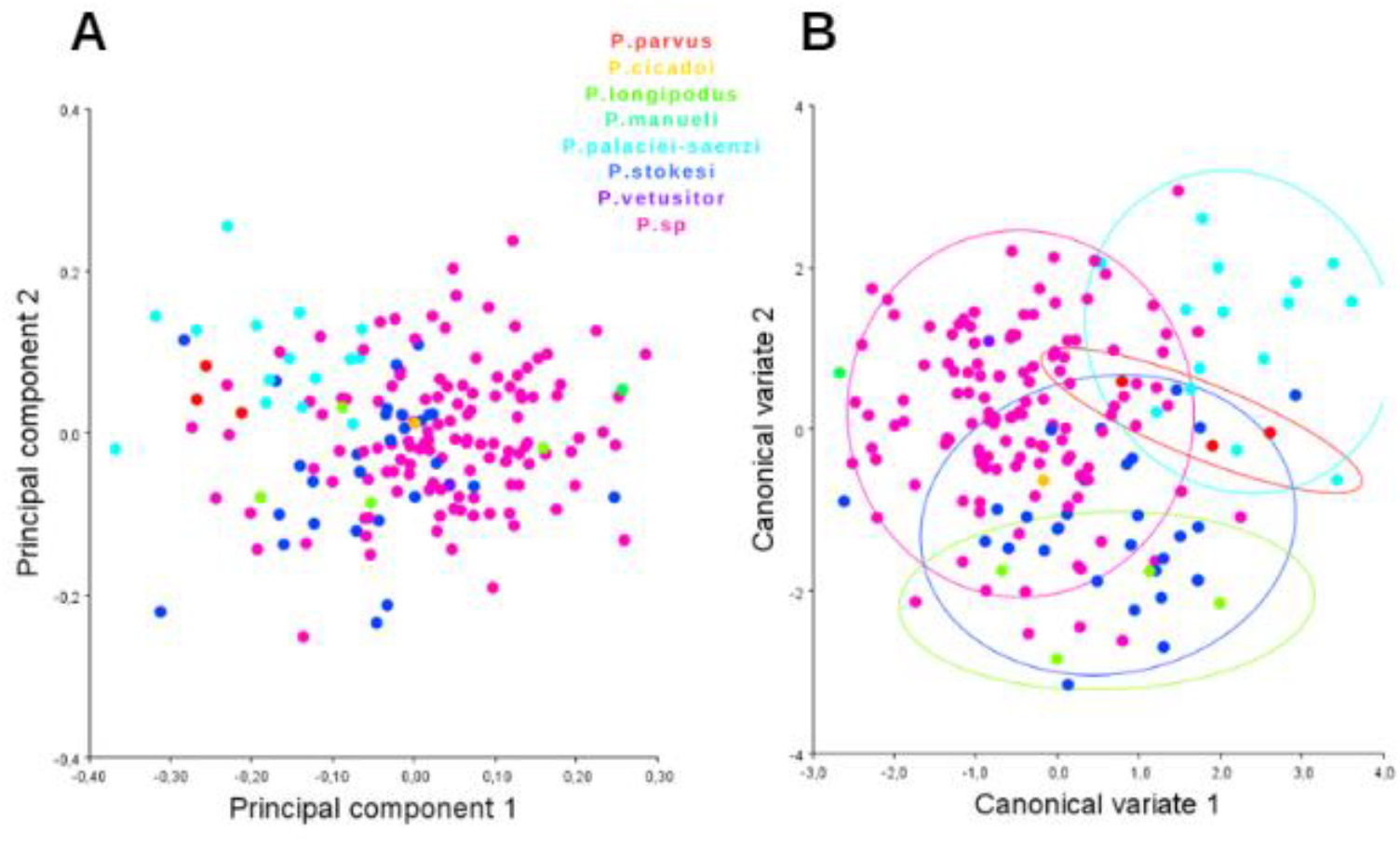
**A**, PCA of the data from the national registry according to the proposed ichnospecies. **B**, CVA following the same criterion (confidence ellipses 0.9 probability).

The result of the regression (Fig. 7) shows a slight variance (2.6%) (p value= 0.0012), suggesting that a small portion of the variance is probably underlined by allometry (ie., size), with larger tracks having wider digits, noted specially in digit I. The remaining 97.4% of the variance is explained by the differences in shape between the footprints analogue as to what wasn’t related to size.

**Figure 7.**
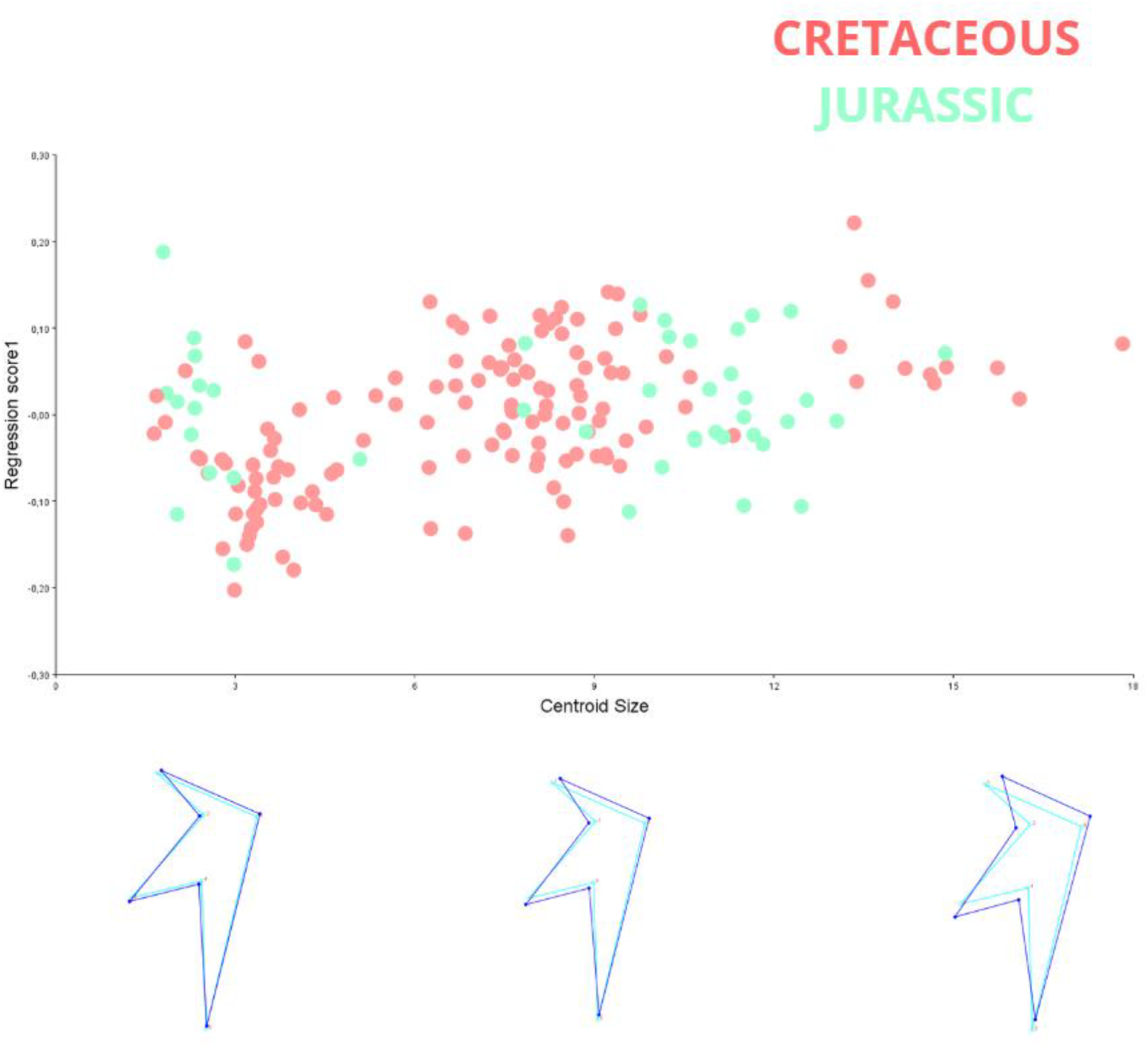
Regression of the national data (Procrustes coordinates as the dependent variable and the size of the centroid as the independent variable) and the diagram of the variation in morphology according to size, compared to the wireframe (mean of all footprints).

## Discussion

The ichnogenus *Pteraichnus* (Stokes, 1957) encompasses many fossil tracks, resulting from the terrestrial locomotion of various groups of pterosaurs. Thus far, the characterization and ichnotaxonomic classification of *Pteraichnus* (as well as the other two proposed ichnogenera: *Purbeckopus* and *Haenamichnus*), has been carried out using the classic tools of paleoichnology, based on the description and comparison of different morphological characters and the measurement of variables such as length, width, and angles between digits. The Iberian pterosaur tracks used in this study, corresponding with the ichnogenus *Pteraichnus*, show a fair continuous variation and distribution thereof, as no clear groupings or significant morphological variations can be ascertained. Although this does not mean that all footprints are identical, clearcut differences cannot be addressed to any of the studied factors, namely, geological time, geographical location or different ichnotaxon. Such homogeneous variability is not surprising, as it is one of paleoichnology’s nemesis, indeed, as it stems from several entwined factors involved in the final track morphology, such as the type of substrate, animal dynamics and, the taphonomic and erosive processes that distort morphology in multiple—and unpredictable—ways (Moratalla et al., 1997; Gatesy & Falkingham, 2017). All in all, we must consider the influence of taphonomic bias, especially when analyzing allometry since our results show a very subtle influence of the size in the shape of the ichnites.

When the age of the ichnites is included in the sample, a continuous distribution is also observed (Fig. 4), even considering a difference of around 90 million years between the oldest and youngest prints. This underscores that the anatomy of pterosaur hands shows little variability over time (referring to digits I, II and III), being a highly conservative character from a morphological point of view. The distribution of the sample according to geographical criteria also shows a very continuous pattern (Fig. 5), in both the tracks from the Spanish record and those from the worldwide record, even if they are distant geographically located or belong to vastly different ecosystems (coastal and/or lacustrine). This further substantiates the fact that it is extremely difficult to classify pterosaur tracks, much as the classification according to the ichnospecies proposed in the literature (seven ichnospecies cited for the Cameros Basin; Fig. 6), which deems it impossible to justify the existence of so many ichnotaxa.

From a biological point of view, our results clearly stress that the morphological homogeneity of pterosaur hands, either both from different clades and from different time across the Mesozoic. Interestingly, although digits I, II and III do not seem to encompass a clear function during flight, they should do in quadrupedal locomotion, as they touch on the sediment and support the anterior proportional part of the animal’s weight. Consequently, the pterosaur track record seems to suggest a strong conservatism (and high morphological convergence) of these structures across pterosaur clades, highlighting the importance of studying the potential selective aspects making these anatomical elements so homogeneous and morphological constancy through pterosaur phylogeny.

## Conclusions

We proposed the study of the ichnological record of the pterosaur manus using shape analysis (Geometric Morphometrics) to mathematically test the extent to which qualitative proxies adequately address pterosaur ichnite variation. The obtained results according to different criteria of classification (temporal, paleogeographic and parataxonomic) have highlighted that the manus tracks of pterosaurs (at least within the ichnogenus *Pteraichnus*), have a very stable morphology, making it extremely difficult to differentiate them attending to any of parataxonomic, palaeogeographic and deep-time age classifiers.

While the obtained results call for the need to revisit such classifications, and a more profound assessment of ichnite taphonomy, the strong resemblance across manus prints proves the conservation of the morphology of their hands. However, rather than flight, the fact that manus anatomy may be important in land locomotion suggests that such a conservative pattern of the organization of the digits I, II and III, may be selective for the pterosaur arm as an adequate element for quadrupedal locomotion.

